# *Structural analysis of Toxoplasma gondii* sortilin

**DOI:** 10.1101/2022.11.17.516902

**Authors:** Ariane Honfozo, Rania Ghouil, Tchilabalo Dilezitoko Alayi, Malika Ouldali, Ana-Andreea Arteni, Cynthia Menonve Atindehou, Lucie Ayi Fanou, Yetrib Hathout, Sophie Zinn-Justin, Stanislas Tomavo

## Abstract

Rhoptries and micronemes are essential for host cell invasion and survival of all apicomplexan parasites, which are composed of numerous obligate intracellular protozoan pathogens including *Plasmodium falciparum* (malaria) and *Toxoplasma gondii* (toxoplasmosis) that infect humans and animals causing severe diseases. We identified *Toxoplasma gondii* TgSORT as an essential cargo receptor, which drives the transport of rhoptry (ROP) and microneme (MIC) proteins to ensure the biogenesis of these secretory organelles. The luminal ectodomain of 752 amino acid long situated at the N-terminus end of TgSORT has been described to bind to MIC and ROP proteins. Here, we present an optimized protocol for expression of the entire luminal ectodomain of TgSORT (Tg-NSORT) in the yeast *Pichia pastoris.* Optimization of its coding sequence, cloning and transformation of the yeast *P. pastoris* allowed the secretion of Tg-NSORT. The protein was purified and further analyzed by negative staining electron microscopy. In addition, molecular modeling using AlphaFold identified key differences between human and *T gondii* sortilin. The structural features that are only present in *T. gondii* and other apicomplexan parasites were highlighted. Elucidating the roles of these specific structural features may be useful for designing new therapeutic agents against apicomplexan parasites

## INTRODUCTION

*Toxoplasma gondii* is a single-celled obligate intracellular parasite responsible for toxoplasmosis. It is the leading cause of congenital neurological abnormalities and severe opportunistic infection in immunocompromised individuals (Hu et al., 2006). This parasite belongs to the large phylum of *Apicomplexa,* which includes protozoan pathogens such as *Babesia, Cryptosporidium, Cyclospora, Isospora* and *Plasmodium* (Kim and Weiss, 2004). Apicomplexan parasites possess various similar morphological features that constitute the hallmark of the phylum (Morrissette and Sibley, 2002). Among these different structures, the most remarkable is the presence of an apical complex composed of polar ring, conoid and unique secretory organelles named rhoptries and micronemes. During the invasion, micronemes firstly release their contents that are required for motility, host cell attachment and egress (Dubois and Soldati-Favre, 2019). Microneme (MIC) proteins (adhesins and escorters) form complexes that link host cell receptors to the glideosome (Opitz and Soldati, 2002). These complexes required for invasion are translocated backwards, allowing the parasite to propel itself into the host cell by membrane invagination. Subsequent proteolysis of these proteins is essential for cell invasion progression by enabling parasite migration into host cell (Brydges et al., 2006). MIC5 is the first microneme protein identified as not possessing an adhesive domain (Brydges et al., 2000). It acts as a regulator of the activity of the parasitic surface protease MPP2 (Brydges et al., 2006). Rhoptries (ROP) proteins are also involved in host cell entry and in the subversion of other host functions such as the control of nuclear transcription and immune responses (Boothroyd and Dubremetz, 2008). Among these proteins, ROP18 phosphorylates and inactivates a family of host immunity related GTPases (IRGs), preserving the parasite from lysis (Fentress and Sibley, 2011). ROP16 interferes with signal transduction in the host nucleus through phosphorylation of the activator STAT3/6 (Butcher et al., 2011). Key roles of rhoptries and micronemes make these organelles the pillars of the parasite survival. However, the formation of these vital secretory organelles depends on the presence of *Toxoplasma gondii* Sortilin-like receptor (TgSORTLR), which is a type I transmembrane cargo transporter located in the post-Golgi and endosome related compartments. TgSORTLR is essential to the biogenesis of secretory organelles and in turn to motility, invasion and egress (Sloves et al., 2012). Furthermore, TgSORTLR is required for efficient host immune responses against infection (Sloves et al., 2015). The luminal ectodomain situated at the N-terminus of the receptor binds ROP and MIC proteins (Sloves et al., 2012) while the cytosolic tail recruits partners to enable anterograde and retrograde receptor transport (Sangaré et al. 2016), in a manner similar to sortilins of humans (Nielsen et al., 2001). Therefore, we now named this receptor TgSORT. Furthermore, homologues of TgSORT is present in all apicomplexan parasites whose genomes have been sequenced (<u>VEuPathDB</u>). For example, in the malaria *P. falciparum,* PfSORT determines transport of proteins to form rhoptries (Hallée et al., 2018a, 2018b).

In humans, sortilins are multifunctional receptors whose structures are defined by the backbone of yeast VPS10 composed of ten β-propeller domains, two cystein bound domains, one transmembrane domain and C-terminus tail (Quistgaard et al., 2009). These sortilins function in mannose-6-phosphate independent manner for the sorting of numerous enzymes from the endosomal system to the lysosomes. Expressed in a number of vertebrate tissues, notably brain, spinal cord, testis, and skeletal muscle, sortilins also function as a surface coreceptor for induction of neural apoptosis in the brain (Kim and Hempstead, 2009) and is linked to type 2 diabetes (Clee et al., 2006) and Alzheimer’s disease (Rogaeva et al., 2007). However, mice or yeast deficient in sortilin/VPS10 are viable and show relatively mild phenotypes (Marcusson et al., 1994; Jansen et al., 2007). In sharp contrast, in apicomplexan parasites, as demonstrated for *T. gondii* and *P falciparum*, this receptor is an essential factor that allows apicomplexan parasites to build the complex apical structure composed of the functional conoid containing rhoptries and micronemes (Sloves et al., 2012; Sakura et al., 2016; Sangaré et al., 2016; Hallée et al., 2018a, 2018b). However, the homology between *T. gondii* sortilin and its human counterparts is only about 27%. Interestingly, four peptide insertions are present in the β-propeller domains B, H, 10CCb and the C-terminus tail in all apicomplexan parasite’s sortilins identified up-to-date (Sloves et al., 2012). The presence of conserved specific peptide insertions exclusively in sortilins of apicomplexan parasites suggests that they may have peculiar 3-D structures compared to the classical VPS10 backbone that can be exploited for future therapeutic interventions. In the present study, we established optimal conditions for expression in the methylotrophic yeast *Pichia pastoris* of secreted soluble N-terminus luminal end of TgSORT for structural analysis and we exploited the recent and powerful computational predictive program AlphaFold to propose a 3D-model of TgSORT.

## MATERIALS AND METHODS

### Strains, vectors and reagents

Pichiapink secreted protein kit (Thermofisher scientific) containing one shot electro-competent *E. coli,* pPINKαHC vector, 5’α-factor primer, *3’CYC1* primer was used in this study. PichiaPink expression strains composed of 4 adenine auxotrophic *Pichia pastoris* strains were as follows: PichiaPink strain 1, a wild strain of genotype *ade2,* PichiaPink strain 2 *(ade2, pep4),* PichiaPink strain 3 *(ade2, prb1)* and PichiaPink strain 4 *(ade2, pep4, prb1).* Pichia Pink Media kit (Dextrose, Pichia Adenine Dropout Agar, yeast extract peptone dextrose, yeast extract peptone dextrose sorbitol, yeast extract peptone dextrose). Additional reagents were required: Yeast Nitrogen Base (Thermofischer scientific), methanol (Millipore), biotin and sorbitol (Sigma Aldrich), glycerol, 1M potassium phosphate buffer. Rabbit anti-Flag antibodies (Sigma Aldrich) and rat anti-Tg-NSORT antibodies were produced in the laboratory as previously described (Sloves et al., 2012). The following media were also prepared: YPD broth, YPD agar, YPDS yeast extract peptone dextrose sorbitol, PAD agar (Pichia Adenine Dropout Agar), BMGY and BMMY.

### *Pichia pastoris* expression vector

The N-terminus luminal coding region of TgSORT (named here Tg-NSORT) from amino acid 37 to 789 (TgSORTLR ToxoDB accession Number TGME49_290160) was designed with Flag and 6xHistidine epitopes added to the C-terminus end. The synthesis and cloning of this Flag and 6xHis tagged TgSORT coding nucleotide sequence in pPINKαHC were performed by Genscript based on *P. pastoris* codon usage. This vector also contained the *Saccharomyces cerevisiae* α-factor signal peptide added upstream to Tg-NSORT (N-terminus) and this allows the traffic of expressed protein to the secretory pathway of the yeast until its release in the culture supernatant. The expression was under the control of the methanol-inducible promoter AOX1.

### Transformation of *P. pastoris*

Strains transformation was made according to Thermofisher scientific recommendations, which are based on modified protocols previously described (Wu and Letchworth, 2004). All cultures were carried out at 27°C, 300 rpm and centrifugations at 1500xg for 5 minutes. The different strains of *Pichia pastoris* were grown on YPD agar plates for 24 hours. Starter cultures were performed by incubating an isolated colony in 10 ml of YPD medium for 24 hours. Hundred ml of cultures were made from an OD_600_ of 0.2 to OD_600_ between 1.3 and 1.5. Pellet was recovered, washed twice in cold sterile water and then in ice-cold 1M sorbitol. Cells were permeabilized for 30 minutes at room temperature using a buffer composed of: 100 mM lithium acetate, 10 mM DTT, 0.6 M sorbitol and 10 mM Tris-HCl pH 7.5. Three washes were performed with ice-cold 1M sorbitol and electro-competent yeast strains were transferred to 0.2-cm electroporation cuvette and 1 μg of linearized recombinant plasmid was added, and kept on ice for 5 minutes. Transformation was performed at 1500 V, 186 Ω and 25 μF using the BTX Electro Cell Manipulator 600. One ml of ice cold YPDS medium was immediately added and the mixture was kept for 2 hours at 27°C without shaking. Ten μl and 100 μl were spread on PAD plates and incubated at 27°C for 24-48 hours until formation of colonies. Positive clones were isolated and cultured for protein expression and purification as described below.

### Recombinant Tg-NSORT protein expression

Protein expression was achieved according to the manufacturer’s protocol. All incubations were done at 27°C under 250 rpm shaking. Pilot experiments were first performed to determine the optimal expression conditions. Ten ml of culture were carried out in BMGY for 24 hours in a 250 ml flask. Cultures were centrifuged at 1500 g for 5 minutes at room temperature and the pellet was resuspended in 1 ml of BMMY. Cells were again incubated overnight before starting the inductions. The four different strains have been tested as well as a range of concentrations of 0.5%, 1%, 2%, 3%, 4% and induction times of 6, 24, 48, 72 and 96 hours. At the end of induction periods, all supernatants were recovered by centrifugation for 10 minutes, analysed by SDS-PAGE and Western blot using the rat anti Tg-NSORT antibodies. *Pichia pastoris* strain transformed with empty vector served as negative control.

After determining the best conditions for expression, a large-scale production was performed. Briefly, one liter of BMGY (Buffered Glycerol-complex Medium) culture medium was seeded by 25 ml using 24-hour pre-culture of one positive clones isolated above. When the OD reaches between 2 and 6, the culture was centrifuged and the pellet was resuspended in 200 ml BMMY (Buffered Methanol-complex Medium) induction medium. After one day of incubation, inductions were performed every 24 hours. Supernatants were analyzed by SDS-PAGE and Western blots. We have also isolated Tg-NSORT from 3 liter of cultures.

### Affinity column purification of Tg-NSORT

200 ml of supernatant containing recombinant Tg-NSORT were concentrated to 5 ml using the 30-kDa cutoff Millipore centrifugal filters. The concentrated sample was diluted 1:10 with 1X binding buffer (50 mM Tris pH 7.5, 250 mM NaCl and 5% glycerol) and then incubated on Nickel-NTA beads for 4 hours at 4°C with 1 mM PMSF and inhibitor cocktail. Three washes were performed in 50 mM Tris pH 7.5, 1 M NaCl and 5% glycerol buffer and three additional washes were done using 50 mM Tris pH 7.5, 150 mM NaCl and 5% glycerol. Recombinant Tg-NSORT was eluted twice with 200 mM of imidazole and eluates were concentrated in 1X PBS, 5% glycerol before size exclusion chromatography. Protein bands were cut and analyzed by mass spectrometry.

### SDS-PAGE

Sixteen (16) μl of supernatant were mixed with 4 μl of 5X Laemmli buffer and heated at 100°C for five minutes. SDS-PAGE was performed in 12% gel under reducing conditions by sample migration at 30 V until its reached running gel then at 70 V and 120 V. Gels were stained with BIO-RAD Coomassie brilliant blue R-250 staining solution. The protein bands of interest were excised and processed for mass spectrometry.

### In-gel digestion of protein and LC-MS/MS analysis

The gel band were cut in small pieces of one mm^3^. The staining of gel pieces were removed thrice with 120 μL of a mixture of 50/50 (v/v) of 25 mM ammonium bicarbonate (NH5CO3)/ acetonitrile for 10 min. In-gel reduction and alkylation of protein disulfide bonds were performed, respectively, with 100 μL of 10 mM of DTT for 50 min at 57 °C, and 100 μL of 50 mM of iodoacetamide (IAM) for 30 min at room temperature. After a washing step with 120 μL of 25 mM NH5CO3 and the dehydration step with 100 μL acetonitrile for 5 min, an in-gel digestion was performed on each band with 0.3 μg of Pierce™ Trypsin Protease, MS Grade (Thermo Fisher Scientific, lL, USA) for 16 h at 37 °C using Thermomixer C (Eppendorf AG, Hamburg, Germany). The peptide was extracted thrice from gel with a mixture of 60/40/0.1 (v/v/v), acetonitrile/25 mM of NH5CO3 (v/v) and 0.1 % formic acid. The extracted solution was then dried with vacuum centrifuge and resuspended in 15 μL of water containing 0.1% formic acid. Seven microliter of each sample were injected into the Ultimate 3,000 RSLC nano-System (Dionex, Thermo Scientific) through a trap column 2 cm x 75 μm inner diameter, C18, 3 μm, 100 A (Dionex, CA, USA) at 3.5 μL/min with aqueous solution containing 0.1% FA and 2% ACN (v/v). After 10 min, the trap column was set on-line with analytical column, EASY-Spray Acclaim PepMap RSLC, 15 cm x 75 μm inner diameter, C18, 2 μm, 100 A (Dionex, CA, USA). Peptides were eluted by applying a mixture of solvents A and B. Solvent A consisted of HPLC grade water with 0.1% FA (v/v), and solvent B consisted of HPLC grade acetonitrile (80% ACN) with 0.1% FA (v/v). Separations were performed using a linear gradient of 2% to 50% solvent B at 300 nL/min over 36 min followed by 3 min linear increase of ACN percentage up to a washing step (4 min at 100% solvent B), 4 min linear decrease of ACN percentage up to an equilibration step (11 min at 2% solvent B). Total analysis run time was 60 min. LC-MS/MS data dependent acquisition was performed using a Q-Exactive HF mass spectrometer (Thermo Scientific, Bremen, Germany) in positive mode. For ionization, an EASY-Spray ES233 (Thermo Scientific, Bremen, Germany) was used with a voltage set at 2 kV, and the capillary temperature set at 350 °C. Full MS scans were acquired in the Orbitrap mass analyzer over an m/z 375 - 1800 range with a resolution set at 60,000 for m/z 200. The target automatic gain control value of 3×10^6^ was used with a maximum allowed injection time (Maximum IT) of 90 ms. For MS/MS, an isolation window of 1.2 m/z was utilized. The ten most intense peaks with a charge state between 2 and 4 were selected for fragmentation using high-energy collision induced dissociation with stepped normalized collision energy of 27-32. The tandem mass spectra were acquired with fixed first mass of m/z 90 in the Orbitrap mass analyzer with a resolution set at 30,000 for m/z 200 and an automatic gain control of 10^4^ The ion intensity selection threshold was 8.3x 10^4^, and the maximum injection time was 120 ms. The dynamic exclusion time was 15 s for the total run time of 60 min.

All data files collected were processed with a specific workflow designed in Proteome Discoverer 2.2 (Thermo Fisher Scientific). MS/MS data was interpreted using Sequest HT search engine (Thermo Fisher Scientific). Searches were performed against *T. gondii* (TGVEG, TGME49 and TGGT1 strains) protein sequences downloaded from www.toxodb.org the 13th November 2019 concatenated with human keratin and others proteins known as contaminants (25310 entries). The search was performed with precursor and fragment mass tolerance respectively set at ±10 ppm and ±0.05 Da and the following dynamic modifications: carbamidomethyl on cysteine, acetyl on protein N-terminal, oxidation on methionine. The target-decoy database search allowed us to control and to estimate the false positive discovery rate at 1% for peptide and protein as well (Elias and Gygi, 2007).

### Western blots

After SDS-PAGE, proteins were transferred on nitrocellulose membrane at 80 V for 1 hour. Membranes were blocked for 30 minutes at room temperature in 5% skim milk prepared in TNT (15 mM Tris-HCl pH8, 140 mM NaCl, 0.05% Tween-20). Incubation with antibodies was done for 1hour at room temperatures with 10 minutes washes for three times using TNT. Blotting membranes were developed with standard chemiluminescent solution (GE healthcare) and scanned using Vilber fusion FX 6.0 apparatus (France).

### Size-exclusion chromatography of Tg-NSORT

Size-Exclusion Chromatography (SEC) using a 24 ml Superose 6 Increase 10/300 GL column in a buffer composed of 50 mM Tris-HCl pH 8, 150 mM NaCl and 1 mM EDTA, was used to purify the concentrated eluate of Tg-NSORT. A single peak corresponding to Tg-NSORT was observed at 17.5 ml by monitoring elution at 280 nm.

### Negative staining of TgN-SORT

Samples were analysed by conventional electron microscopy using the negative staining method. 3 μL suspension (0.05 mg mL^-1^) were deposited on an airglow-discharged carbon-coated grid. Excess liquid was blotted, and the grid rinsed with 2% w/v aqueous uranyl acetate. The grids were visualised at 100 kV with a TECNAI Spirit (FEI) transmission electron microscope (ThermoFisher, New York NY, USA) equipped with a K2 Base 4k × 4k camera (Gatan, Pleasanton CA, USA). Final magnification was at 34.500 x, corresponding to a pixel size at the level of the specimen of 0.14 nm. Data were recorded under low-dose conditions (dose rate 20 e A^-2^).

## RESULTS

### Determination of optimal conditions for Tg-NSORT expression

We have used four distinct yeast *Pichia pastoris,* which were mutants lacking respectively either one gene coding a first protease, a second protease or a double mutant lacking both proteases and wild type strains. The use of these mutants allows to minimize the rate of degradation of the recombinant protein expressed in the yeast and to achieve an optimal expression in one of these four *P. pastoris* strains. Figure 1A depicted the plasmid that contains the coding DNA sequence of Tg-NSORT tagged to FLAG and 6XHis epitopes and used to transform these yeasts. Transformation of these *P. pastoris* strains resulted in good integration of the plasmids into the genome. Two positive colonies of each strain were shown to express a protein having about 100 kDa band that corresponds to the expected size of Tg-NSORT (Figure 1B). However, different amounts of the 100-kDa protein were observed after Coomassie blue staining plus few additional smaller bands with one predominant band around 50 kDa (Figure 1B). We showed that the specific anti-TgSORT antibodies recognized this 50-kDa protein, suggesting it as a degradation product of the apparently intact and much stronger TgN-SORT band of 100-kDa size (Figure 1C). Based on the level and intactness of TgN-SORT, we selected clone 1 of *Pichia pastoris* mutant strain 4 for our studies (Figure 1B and 1C, see the red box). Using this Tg-NSORT clone 1, we established that the same level of the 100-kDa band was expressed regardless of the methanol concentrations, except that a slight decreased in intensity was observed at 4% methanol (Figure 1D and 1E). In addition, the amount of TgN-SORT produced in this clone also increased with time (Figure 1F and 1G). For our purposes, we picked out 2% of methanol induction for 72 hours as the optimal conditions at 27°C under constant shaking for efficient expression of Tg-NSORT in *Pichia pastoris.*

**Figure 1.**
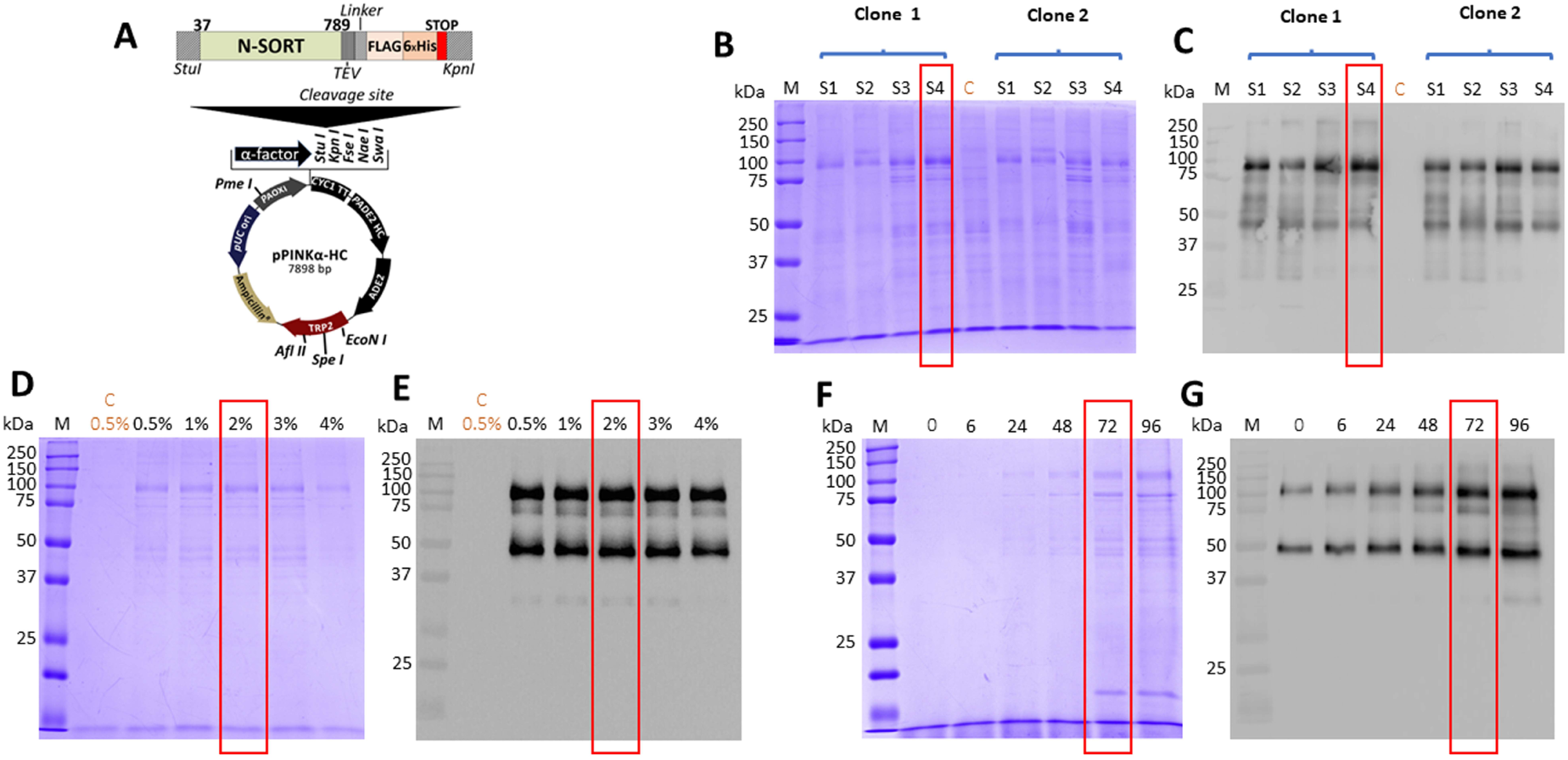
(A) Design of the expression vector of TgSORT in *Pichia pastoris.* (B) A pilot experiment of Tg-NSORT expression and transformants derived from *Pichia pastoris* S1, S2, S3 and S4-strains were analysed by SDS-PAGE and Coomassie blue staining. (C) Western blot of this pilot experiment using anti-TgSORT antibodies; (D) The optimal methanol concentration for induction determined by Coomassie bue staining; (E) The corresponding blots of panel D; (E) Coomassie blue staining showing the optimal time (hours) for methanol induction; (F) The corresponding blot of panel E. C means negative control.

### Purification of secreted Tg-NSORT from *Pichia pastoris*

After, a small-scale purification of Tg-NSORT using Ni-NTA resin, we checked that the 100 kDa and 50 kDa bands were recognized by Western blots using anti-His and anti-Flag antibodies corresponding to the two-epitope tags placed at the C-terminus of Tg-NSORT (Figure 2A). The same bands stained by Coomassie blue after polyacrylamide gel electrophoresis were excised and processed for mass spectrometry. Sixty-three (63) peptides covering 55% of the length of TgSORT were identified in the 100-kDa protein, indicating it is a genuine ectodomain of Tg-NSORT expressed and secreted from *P. pastoris* (panel B of Figure 2). The nature of these peptides was described in Table 1, which also showed the sequence of 46 peptides found in the 50-kDa protein, suggesting it as a degraded product of Tg-NSORT. Altogether, these data demonstrate that Tg-NSORT was expressed and secreted by *P. pastoris.* After this verification, we embarked on a large-scale production and purification of Tg-NSORT. Figure 3 shows the quality of Tg-NSORT secreted by *P. pastoris* in one liter of culture medium, which was used to purify the protein by Ni^+^-NTA beads (Figure 3A and 3B). The highest amount of protein was obtained after 200 mM of imidazole elution (see E2) and this yielded to 0.5 mg of total Tg-NSORT protein with fewer degradation (Figure 3A and 3B). The increase of the volume of *P. pastoris* culture to three liters resulted in a higher quantity of protein that reached 2 mg of purified protein (Figure 3B and 3D). Next, we decided to improve the purity of Tg-NSORT by removing the smaller degraded products seen in Figure 3 by size exclusion chromatography. In these gel filtration conditions, Tg-NSORT was eluted at 17.5 ml, which corresponds to a molecular mass between 44 and 158 kDa on this column (Figure 4A). SDS-PAGE revealed Tg-NSORT at about 100 kDa, but the degraded product of 50 kDa was still present after gel filtration, suggesting that it binds to the 100 kDa protein (Figure 4B).

**Figure 2.**
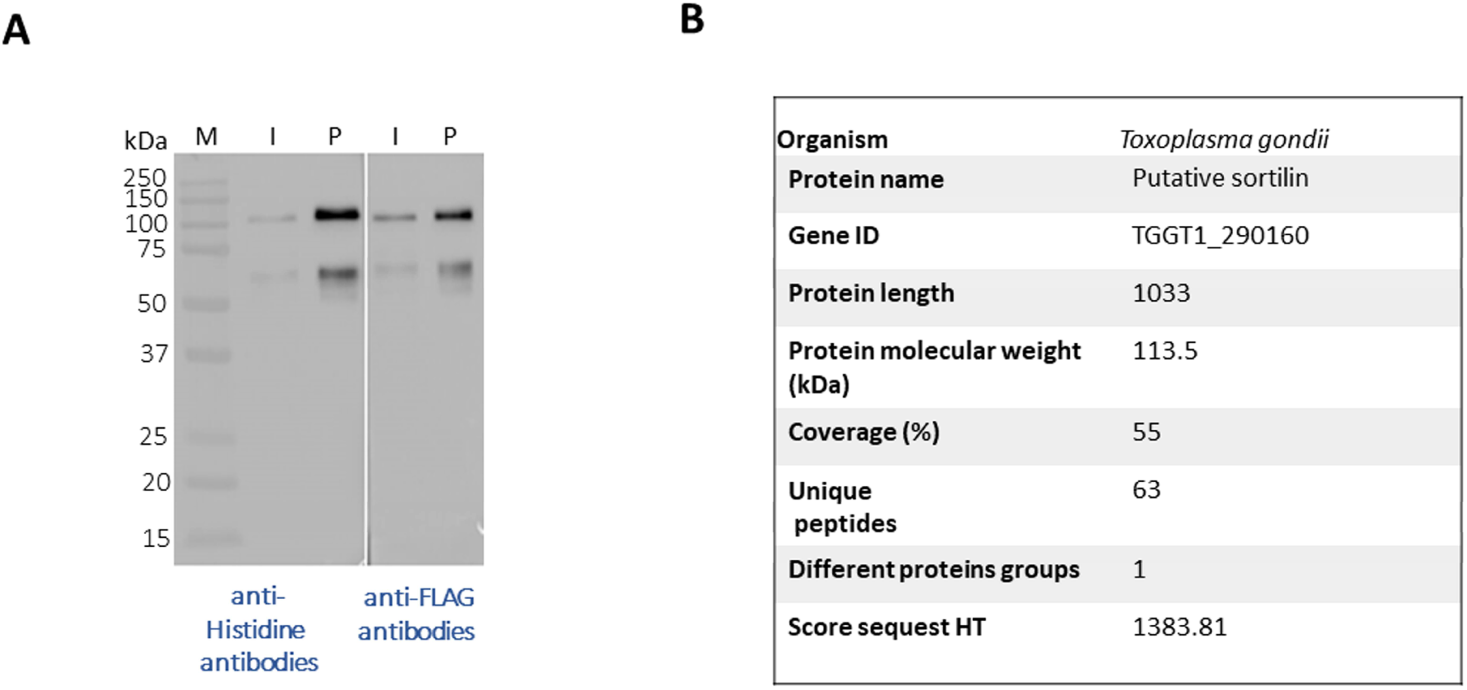
(A) Western blots after a small-scale purification of Tg-NSORT using Ni^+^NTA column and the culture supernatant of *Pichia pastoris* induced by 2% of methanol for 3 days. The blots were revealed with anti-Histidine and anti-Flag, two epitope tags at the C-terminus of Tg-NSORT. (B) The same material analysed in panel A was staining by Coomassie blue and processed by mass spectrometry to confirm the 100-kDa band as Tg-NSORT.

**Figure 3.**
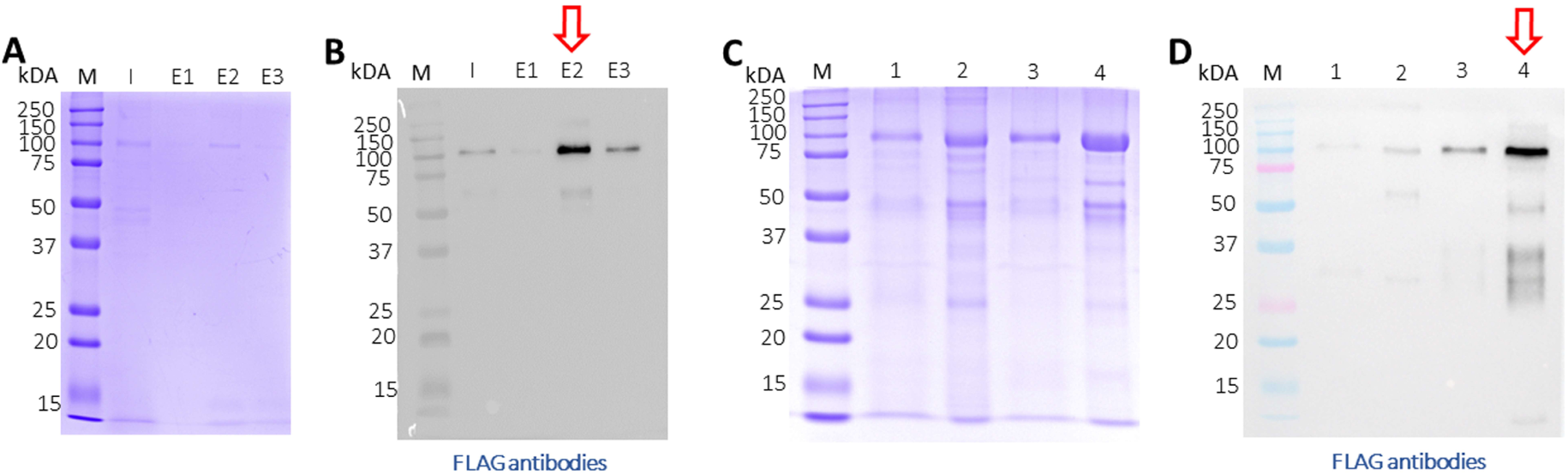
Large-scale purification of TgN-SORT by affinity chromatography (A) Coomassie blue staining and corresponding blot after purification from one liter of st *Pichia pastoris* transformant grown in complete medium culture; I: Input; E1: 1 eluate nd rd at 50 mM of imidazole; E2: 2^nd^ eluate at 200 mM of imidazole; E3: 3 eluate at 200 mM of imidazole; (C and D) the same experiment using three liters of culture medium containing secreted TgSORT by *Pichia pastoris;1:* Input 1X; 2: Input TgN-SORT 20X material; 3: 1X eluate of TgN-SORT; 4: 15X eluate of TgN-SORT.

**Figure 4.**
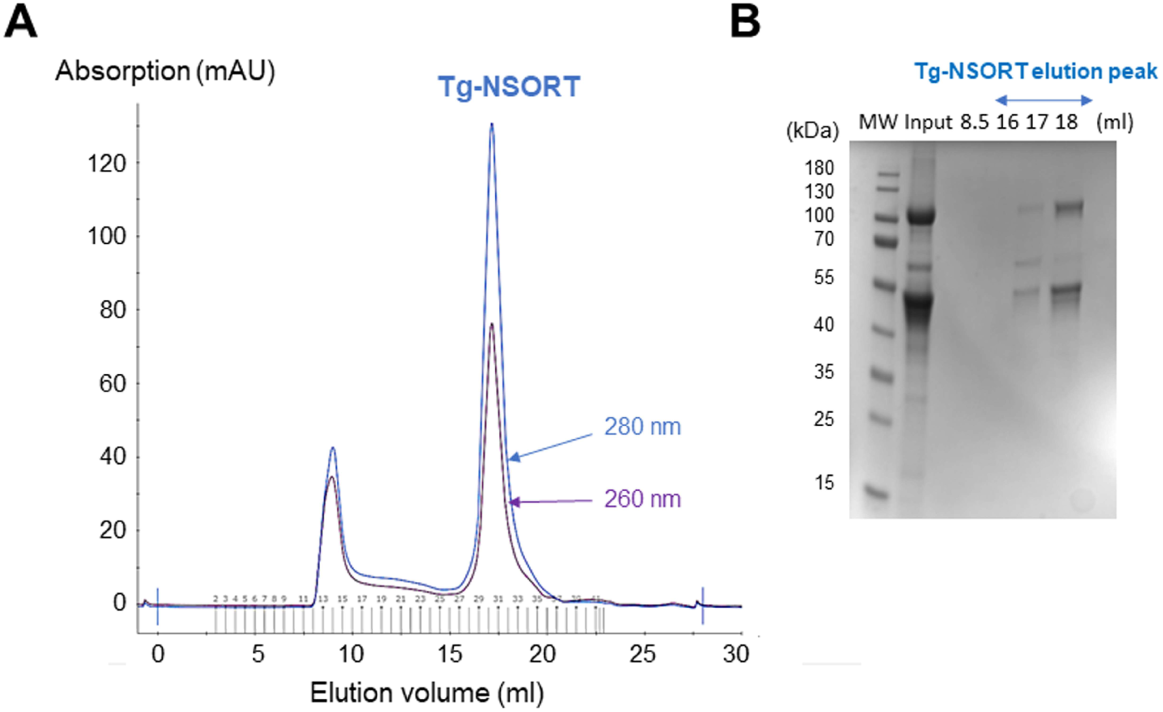
Size exclusion chromatography of Tg-NSORT. (A) Chromatogram recorded at 260 (purple) and 280 (blue) nm showing the peak corresponding to eluted Tg-NSORT. (B) SDS-PAGE and Coomassie blue staining of eluate containing Tg-NSORT.

### Negative staining electron microscopy

We diluted the purified material from size chromatography down to 0.05 mg/ml concentration to analyze it by conventional Electron Microscopy (EM) using the negative staining method. We obtained EM micrographs showing homogeneous and well-dispersed particles, suggesting that a protein having a single conformation was present in our gel filtration samples (Figure 5A). Analysis of these micrographs revealed a ring-shaped protein structure that resembles that previously shown for human sortilin (Figure 5B). However, further attempts to crystallize Tg-NSORT or to analyze it by cryo-electron microscopy failed. Therefore, we used the recent AlphaFold2 program (Jumper et al., 2021) to calculate a model of Tg-NSORT.

**Figure 5.**
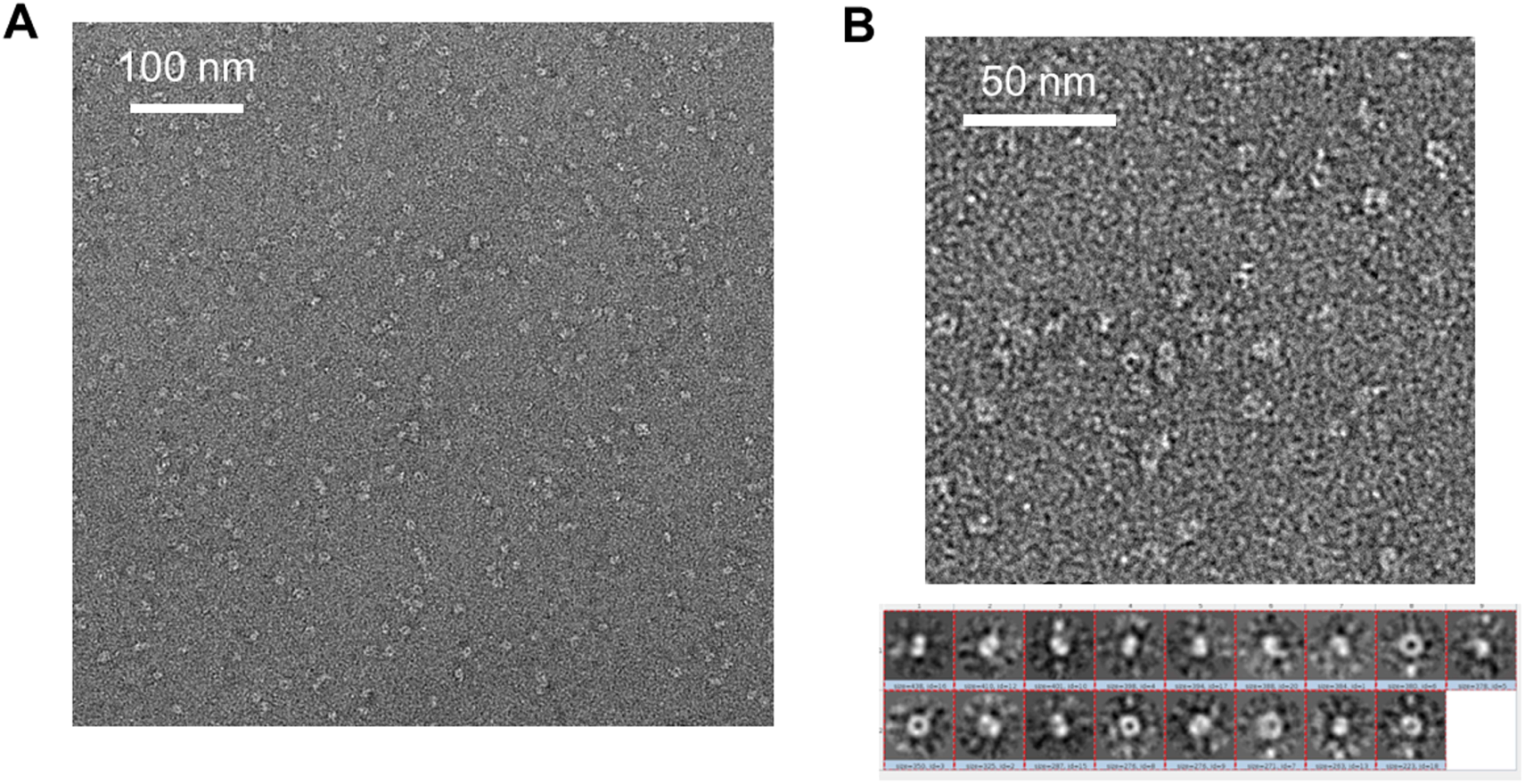
(A) Negative staining electron micrograph of Tg-NSORT purified by gel filtration. (B) Zoom on a micrograph and 2D classification of the particles picked on the micrographs.

### AlphaFold analyses

We provided as an input for AlphaFold2 the sequence of Tg-NSORT. Using this recent program, we obtained a three-dimensional model of Tg-NSORT (Figure 6A), and in particular of the four parasite’s specific peptide insertions, which are organized as loops within TgSORT (Figures 6A and 6C). We also attempted to model the 3D structure of Tg-NSORT bound to its partners. AlphaFold2 predicted with a reasonable significant score (lDDT values for the residues of the disordered binding partners larger than 0.7) that Tg-NSORT interacts with different ROPs proteins (ROP1, ROP5, ROP16) through specific motifs located in intrinsically disordered regions (IDRs) of ROPs proteins (see the model of Tg-NSORT bound to ROP1 in Figure 7A). These motifs are found in the pro-peptide situated at the N-terminus of all ROP proteins. They all bind to the same site in Tg-NSORT within the inner tunnel of the protein. Such binding mode was already observed for neurotensin binding to human sortilin (Figure 7B; Quistgaard et al., 2009).

**Figure 6.**
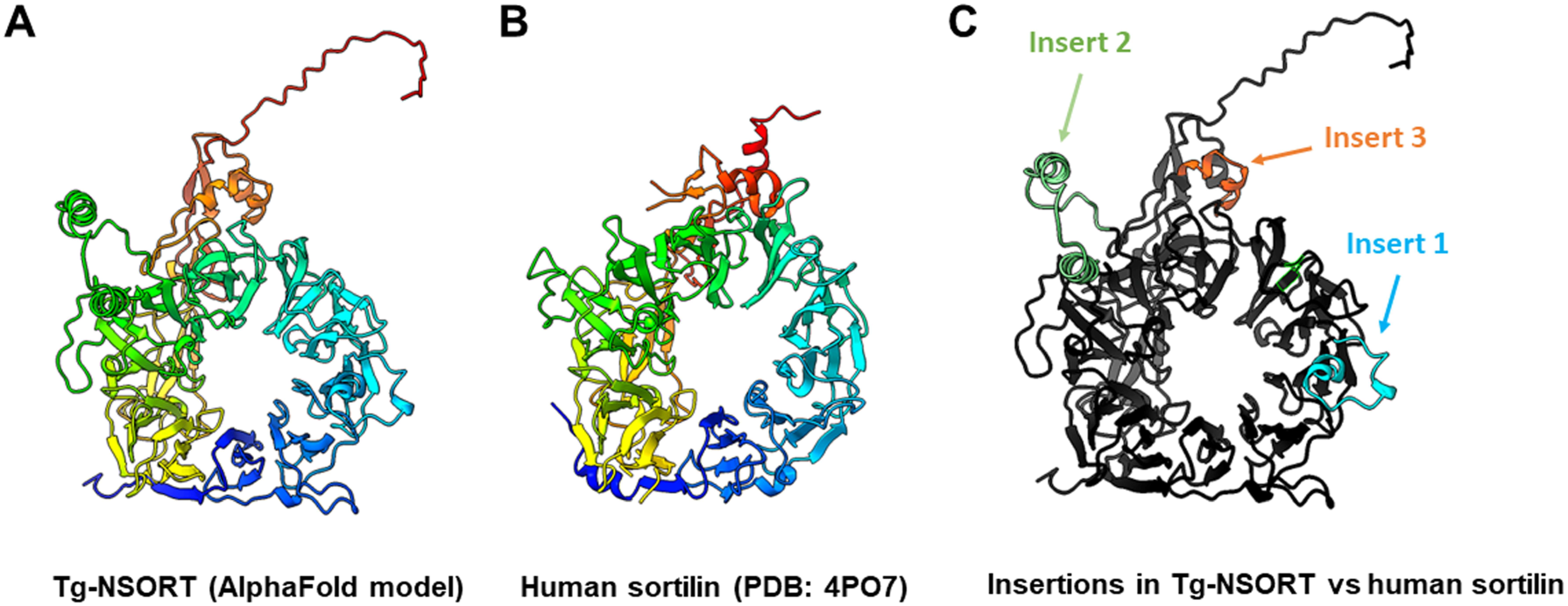
(A) 3D model of Tg-NSORT calculated using AlphaFold2. (B) Crystal structure of human sortilin. (C) Representation of the Tg-NSORT model, with the insertions specific to *Apicomplexa* colored in cyan (insert or loop 1), green (insert or loop 2) and orange (insert or loop 3). In panels (A) and (B), proteins are colored from blue (N-terminus) to red (C-terminus).

**Figure 7.**
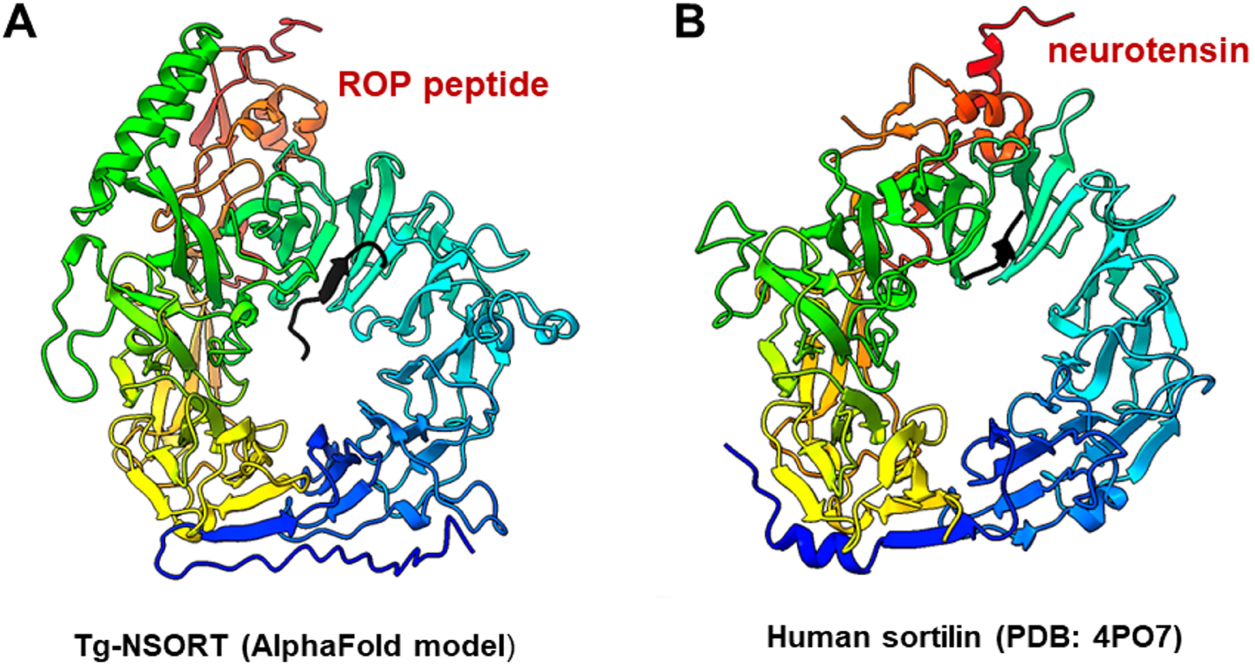
(A) Interaction between Tg-NSORT (from blue to red) and the intrinsically disordered region (IDR) of ROP1 (in black), as determined by AlphaFold2. The motif PPNAQELLPP of this IDR binds to the tunnel formed by the ten β-propeller domains of sortilin. (B) Interaction between human sortilin and the neurotensin peptide, as observed in the crystal structure referenced as 4PO7 in the PDB.

## DISCUSSION

*Pichia pastoris* system has a good record of accomplishment in expressing proteins from both prokaryotes and eukaryotes for the most part difficult to express in *E. coli.* Human coagulation factor XIIIa, which was insoluble after its complicated expression in *E. coli* (Nikolajsen et al., 2014) was successfully produced by Chen et al., (2021) using *P. pastoris*. As used in the present study, these authors also employed the methanol inducible AOX1 promoter to efficiently express the protein for which they evaluated different biological activities (Chen et al., 2021). The same AOX1 promoter controls the expression of the human camel chymosin in *P. pastoris* (Wang et al., 2015). Protein expression in *P. pastoris* depends of different parameters including pH of the culture medium, inducing agent and temperature. As for human coagulation factor XIIIa, the ectodomain of *T. gondii* TgSORT composed of the ten β-propeller and double 10 C-C bound domains is insoluble because it is completely directed into inclusion bodies when expressed in *E. coli*. Now, we have been quite successful in expressing the ectodomain of TgSORT, which is transported through the secretory pathway and can be recovered in the culture supernatant as several milligrams of proteins using *P. pastoris*. As this yeast is a eukaryotic model system; we assume that the parasite TgSORT will be well folded in addition to some post-translational modifications that may be necessary for future functional investigations. It should be noticed that, for example, *T. gondii* ROP2 protein has been already expressed in *P. pastoris* and the protein has been used for diagnosis (Chang et al., 2011). However, the level of ROP2 in *P. pastoris* appears weaker than what we have achieved during our work. After 72 hours of induction with 2% of methanol, we were able to obtain several milligrams of pure recombinant TgSORT. It is known that the experimental conditions are important for protein expression in this yeast. For example, Cheng et al. (2021) induced the expression at 30°C with 1% of methanol every 24 hours during 120 hours for a camel chymosin at pH 4.7 and even below at 28°C for 8 hours (Wang et al., 2015). After evaluation of the different parameters, expression of human serum albumin was performed at 28°C for 24 hours in an acidic pH of 5.75 and the methanol concentration was set between 0.5 and 2% and for every 2 hours (Zhu et al., 2018). In addition, of these conditions defined for human proteins expression in *P. pastoris,* other *T. gondii* proteins whose expression was difficult in *E. coli* were also produced in this yeast. TgGRA2 was expressed in *P. pastoris* at 28°C for 5 days under 0.5% final methanol every 24 hours (Ling et al., 2012), (Huaiyu Zhou, 2007). GRA4 was also expressed in under the same conditions but for 3 days (Lau et al., 2010). In addition to these *T. gondii* cytosolic proteins, membrane proteins such as SAG1 and SAG2 were also expressed in *P. pastoris* at 30°C for 4 days under continuous 24 hours of induction using 0.5-1% methanol concentrations (Thiruvengadam et al., 2011), (Huaiyu Zhou, 2007), (Lau Yee Ling et al., 2010). Methanol concentration generally varies between 0.5% and 2% and even very low amount can stimulate induction between short time of culture as it represents an important source of carbon for the yeast. Its continuous addition in the culture favors the good expression of proteins. In addition, we also noticed that even a minimal expression environment could be achieved if good aeration conditions of the cultures were established. Using *P. pastoris,* we obtained enough quantity of purified Tg-NSORT that was analyzed by negative-staining electron microscopy. The purified protein was homogenous, and we observed ring-shaped particles with dimensions similar to those of the crystal structure of human sortilin (Quistgaard et al., 2009). We were to obtain a higher atomic resolution structure of Tg-NSORT. It seems that the presence of the three parasite’s specific loops hinders crystallization of Tg-NSORT. Alternatively, we used the AlphaFold program for predicting the structure of Tg-NSORT. This analysis revealed the possible molecular bases of the interactions between TgSORT and ROP proteins. It indicated that the propeptide domain of ROP could bind to the tunnel of TgSORT in a manner similar to neurotensin with human sortilin. The specific motifs of ROP proteins are located in some intrinsically disordered regions (IDR) with all IDR tested binding to the same site of TgSORT. These IDR corresponds to the pro-peptide situated at the N-terminus of all ROP proteins, and it is normally cleaved off during the maturation of ROP proteins before they reached their destination (Bradley and Boothroyd, 1999; Bradley and Boothroyd, 2002; et al., Hajagos et al., 2012).

In conclusion, *Pichia pastoris* represents a good organism for *T. gondii* protein expression. This expression system combines the advantages of prokaryotic system with those of eukaryotic model for use of minimal and inexpensive culture medium, fast and high growth rate, high productivity, extracellular expression, folding and post-translational modifications (Karbalei et al., 2020). Large-scale production in improved conditions can lead to high amounts of well-folded proteins that can be used for various applications. Collectively, our data provides the foundation of future and deeper structure-function studies of TgSORT, the key receptor required for host infection of apicomplexan parasites. The expression of large amount of soluble ectodomain of TgSORT provides an avenue for conducting detailed mechanistic studies with biochemical and chemical approaches to identify new and parasite-specific inhibitors.

---

**Supplementary Table 1.** Mass spectrometry data showing peptide identified from the 100-kDA and 50-kDA proteins of purified TgSORT excised from polyacrylamide gel and stained by Coomassie blue and processed for proteomics analyses.

## Acknowledgments

Financial support for this work was provided by the Agence Nationale de la Recherche grants N°ANR-19-CE44-0006. AH was supported by the International Campus France and African UEMOA fellowships. This work benefited from the CryoEM platform of I2BC, supported by the French Infrastructure for Integrated Structural Biology (FRISBI) [ANR-10-INSB-05-05]. We thank Dr. P. Legrand (Synchrotron Soleil) for the AlphaFold2 calculations.

## Competing interest statement

The authors declare that they have no competing financial interest

## Author Contributions

The author(s) have made the following declarations about their contributions:

## Conceived and designed the experiments

SZJ and ST

## Performed the experiments

AH, RG, TDA, AA, MOA and SZJ

## Contributed reagents/materials/analysis tools

SZJ, YA and ST

## Performed data analysis

AH, CMA, LAF, SZJ

## Wrote the paper

AH, SZJ, ST

## Notes

### Competing Interest Statement

The authors have declared no competing interest.

## REFERENCES

Boothroyd, J.C., Dubremetz, J.-F., 2008. Kiss and spit: the dual roles of *Toxoplasma* rhoptries. Nat Rev Microbiol 6, 79–88. https://doi.org/10.1038/nrmicro1800

Bradley, P. J., and Boothroyd, J. C. Identification of the pro-mature processing site of *Toxoplasma* ROP1 by mass spectrometry. Mol. Biochem. Parasitol. 1999.100,103–109.

Bradley, P. J., and Boothroyd, J. C. The pro region of *Toxoplasma* ROP1 is a rhoptry-targeting signal. Intern. J. Parasitol. 2001. 31,1177–1186.

Brydges, S.D., Sherman, G.D., Nockemann, S., Loyens, A., Däubener, W., Dubremetz, J.-F., et al. 2000. Molecular characterization of TgMIC5, a proteolytically processed antigen secreted from the micronemes of *Toxoplasma gondii*. Mol. Biochem. Parasitol. 111, 51–66. https://doi.org/10.1016/S0166-6851(00)00296-6

Brydges, S.D., Zhou, X.W., Huynh, M.-H., Harper, J.M., Mital, J., Adjogble, K.D.Z., et al. Targeted deletion of *MIC5* enhances trimming proteolysis of *Toxoplasma* invasion proteins. Eukaryot Cell 5, 2174–2183. https://doi.org/10.1128/EC.00163-06

Butcher, B.A., Fox, B.A., Rommereim, L.M., Kim, S.G., Maurer, K.J., Yarovinsky, F., et al. 2011. *Toxoplasma gondii* Rhoptry Kinase ROP16 Activates STAT3 and STAT6 Resulting in Cytokine Inhibition and Arginase-1-Dependent Growth Control. PLoS Pathog 7, e1002236. https://doi.org/10.1371/journal.ppat.1002236

Chang, P. Y., Fong, M. Y., Nissapatorn, V., and Lau, Y. L. Evaluation of *Pichia pastoris*- expressed recombinant rhoptry protein 2 of *Toxoplasma gondii* for its application in diagnosis of toxoplasmosis. Am J Trop Med Hyg. 2011. 85, 485–489. doi: 10.4269/ajtmh.2011.11-0351.

Clee, S. M., Yandell, B. S., Schueler, K. M., Rabaglia, M. E., Richards, O. C., Raines, S. M., et al. (2006). Positional cloning of Sorcs1, a type 2 diabetes quantitative trait locus. Nat. Genet. 38, 688–693.

Dubois, D.J., Soldati-Favre, D., 2019. Biogenesis and secretion of micronemes in *Toxoplasma gondii*. Cellular Microbiology 21, e13018. https://doi.org/10.1111/cmi.13018

Elias, J. E., and Gygi, S. P. Target-decoy search strategy for increased confidence in large-scale protein identifications by mass spectrometry. 2007. Nat. Methods 4, 207–214

Fentress, S.J., Sibley, L.D., 2011. The secreted kinase ROP18 defends *Toxoplasma’s* border. BioEssays 33, 693–700. https://doi.org/10.1002/bies.201100054

Hajagos, B. E, Turetzky, J. M., Peng, E. D., Cheng, S. J., Ryan, C. M., Souda, P., et al. Molecular dissection of novel trafficking and processing of the*Toxoplasma gondii* rhoptry metalloprotease toxolysin-1. Traffic. 2012 Feb;13(2):292–304. doi: 10.1111/j.1600-0854.2011.01308

Hallée, S., Boddey, J. A., Cowman, A. F., and Richard, D. Evidence that the *Plasmodium falciparum* protein sortilin potentially acts as an escorter for the trafficking of the rhoptry-associated membrane antigen to the rhoptries. mSphere. 2018a. 3: e00551–17. doi: 10.1128.

Hallée, S., Counihan, N. A., Matthews, K., de Koning-Ward, T. F., and Richard, D. The malaria parasite *Plasmodium falciparum* sortilin is essential for merozoite formation and apical complex biogenesis. Cell Microbiol. 2018b. 20, e12844. doi: 10.1111.

Hu, K., Johnson, J., Florens, L., Fraunholz, M., Suravajjala, S., DiLullo, C., et al. 2006. Cytoskeletal Components of an Invasion Machine-The Apical Complex of *Toxoplasma gondii*. PLoS Pathog. 2, e13. https://doi.org/10.1371/journal.ppat.0020013

Jansen, P., Giehl, K., Nyengaard, J. R., Teng, K., Lioubinski, O., Sjoegaard, S. S., et al. (2007). Roles for the pro-neurotrophin receptor sortilin in neuronal development, aging and brain injury. Nat. Neurosci. 10, 1449–1457.

Jumper, J., Evans, R., Pritzel, A., Green, T., Figurnov, M., Ronneberger, O., et al. Highly accurate protein structure prediction with AlphaFold. Nature. 2021. 596, 583–589. doi: 10.1038/s41586-021-03819-2.

Kim, T., and Hempstead, B. L. (2009). NRH2 is a trafficking switch to regulate sortilin localization and permit proneurotrophin-induced cell death. EMBO J. 28, 1612–1623.

Kim, K., Weiss, L.M., 2004. *Toxoplasma gondii:* the model apicomplexan. Int J Parasitol 34, 423–432. https://doi.org/10.1016/j.ijpara.2003.12.009

Quistgaard, E. M., Madsen, P., Groftehauge, M. K., Nissen, P., Petersen, C. M., and Thirup, S.S. Ligands bind to Sortilin in the tunnel of a ten-bladed beta-propeller domain. Nat. Struct. Mol. Biol. 16, 96–98, (2009)

Marcusson, E. G., Horazdovsky, B. F., Cereghino, J. L., Gharakhanian, E., and Emr, S. D. (1994). The sorting receptor for yeast vacuolar carboxypeptidase Y is encoded by the VPS10 gene. Cell 77, 579–586.

Marti-Renom, M. A., Stuart, A., Fiser, A., Sánchez, R., Melo, F., and Sali, A. Comparative protein structure modeling of genes and genomes. Annu. Rev. Biophys. Biomol. Struct. 29, 291–325, 2000.

Morrissette, N.S., Sibley, L.D., 2002. Cytoskeleton of apicomplexan parasites. Microbiol. Mol. Biol. Rev. 66, 21–38. https://doi.org/10.1128/MMBR.66.1.21-38.2002

Nielsen, M. S., Madsen, P., Christensen, E. L., Nykjaer, A., Gliemann, J., Kasper, D., et al. The sortilin cytoplasmic tail conveys Golgi-endosome transport and binds the VHS domain of the GGA2 sorting protein. EMBO J. 2001 May 1;20(9):2180–90. doi: 10.1093/emboj/20.9.2180.

Nikolajsen, C.L., Dyrlund, T.F., Poulsen, E.T., Enghild, J.J., and Scavenius, C. 2014. Coagulation factor XIIIa substrates in human plasma. J. Biol. Chem. 289, 6526–6534. https://doi.org/10.1074/jbc.M113.517904

Opitz, C., Soldati, D., 2002. ‘The glideosome’: a dynamic complex powering gliding motion and host cell invasion by Toxoplasma gondii. Mol. Microbiol. 45, 597–604. https://doi.org/10.1046/j.1365-2958.2002.03056.x

Sangaré, L. O., Alayi, T. D., Hovasse, A., Westermann, B., Sindikubwabo, F., Callebaut, I., et al. Unconventional endosome-like compartment and retromer complex in *Toxoplasma gondii* govern parasite integrity and host infection. Nat. Commun. 2016. 7:10191 doi: 10.1038/ncomms11191.

Sloves, P.-J., Delhaye, S., Mouveaux, T., Werkmeister, E., Slomianny, C., Hovasse, A., et al. 2012. *Toxoplasma* sortilin-like receptor regulates protein transport and is essential for apical secretory organelle biogenesis and host infection. Cell Host Microbe 11, 515–527. https://doi.org/10.1016/j.chom.2012.03.006

Sloves, P.-J., Mouveaux, T., Ait-Yahia, S., Vorng, H., Everaere, L., Sangare, L. O., al. 2015. Apical organelle secretion by *Toxoplasma* controls innate and adaptive immunity and mediates long-term protection. J. Infect. Dis. 212, 1449–1458. https://doi.org/10.1093/infdis/jiv250

Wu, S., Letchworth, G. J., 2004. High efficiency transformation by electroporation of *Pichia pastoris* pretreated with lithium acetate and dithiothreitol. BioTechniques 36, 152–154. https://doi.org/10.2144/04361DD02.

